# Persistent representation of a prior schema in the orbitofrontal cortex facilitates learning of a conflicting schema

**DOI:** 10.1101/2025.02.28.640679

**Authors:** Ido Maor, James Atwell, Ilana Ascher, Yuan Zhao, Yuji K. Takahashi, Evan Hart, Francisco Pereira, Geoffrey Schoenbaum

## Abstract

Schemas allow efficient behavior in new situations, but reliance on them can impair flexibility when new demands conflict, culminating in psychopathology. Evidence implicates the orbitofrontal cortex (OFC) in deploying schemas in new situations congruent with previously acquired knowledge. But how does this role affect learning of a conflicting behavioral schema? Here we addressed this question by recording single-unit activity in the OFC of rats learning odor problems with identical external information but orthogonal rules governing reward. Consistent with schema formation, OFC representations adapted to track the underlying rules, and both performance and encoding was faster on subsequent than initial problems. Surprisingly however, when the rule governing reward changed, persistent representation of the prior schema was correlated with acquisition of the new. Thus, OFC was not a source of interference and instead supported new learning by accurately and independently representing the old schema as the new was acquired.

## Introduction

Understanding the rules that govern a specific situation and generalizing them to other situations with similar structures is a fundamental cognitive ability essential for adaptive behavior ^1–6^. This process acts as a mental shortcut, enabling efficient problem-solving through the application of preexisting templates constructed from prior experience in new situations ^7–14^. The use of such templates, or schemas, is evident across a wide range of tasks and behaviors, where identifying key rules and extracting abstract principles leads to improved performance over time ^15–19^. For instance, experienced chess players often recognize patterns such as controlling the center, maintaining piece activity, or leveraging pawn structures and use these abstract strategies to guide their decisions across diverse positions.

The OFC is thought to play a pivotal role within a circuit mediating the development and use of behavioral schemas, specializing in the identification of hidden states – rules – that generalize across similar problems ^20–22^. While the contribution of the OFC to this cognitive function is a relatively new proposal, its long-appreciated role supporting rapid reversal learning can be viewed as an example of this function ^23–26^; so can the role of OFC in settings such as devaluation ^27, 28^, in which a new goal value must be generalized to novel situations.

Yet, the application of prior knowledge can introduce incorrect assumptions or biases in new situations governed by conflicting rules. For example, a chess player transitioning to Go might mistakenly apply chess strategies, like controlling the center, which is fundamental in chess but can be ineffective or even counterproductive in Go where the center is often secondary to corners and sides in the early game. Failure to adapt to the appropriate schema in new situations can result in ineffective and even maladaptive behaviors, which are most prominently exemplified in conditions like obsessive-compulsive disorder ^29, 30^ and addiction ^31, 32^.

If the OFC supports the formation and use of schemas, it becomes of interest how it manages orthogonal or conflicting behavioral schemas. One possibility is that the neural representation of a previously learned schema is silenced, overwritten, or jettisoned quickly when a new schema is encountered. Alternatively, the OFC might maintain parallel representations of both old and new schemas, allowing for flexible switching between them as environmental demands shift. This would allow the old schema to remain active to influence behavior but might also hinder or prevent efficient adoption of the new schema.

Here we adjudicated between these possibilities by recording single-unit activity in the OFC of rats during learning of a series of odor problems in which external information was identical, but the rules governing reward were orthogonal. As expected, we found that OFC representations adapted to track the underlying rules, and both performance and encoding were faster on subsequent than initial problems, consistent with schema formation. In rats trained on both rules, the OFC persistently multiplexed information pertaining to the two schemas and, surprisingly, this interleaved representation had a positive effect on the acquisition of the new conflicting schema.

## Results

### Behavioral performance during learning of orthogonal rules

Rats were trained on a series of odor-guided discrimination problems, involving two sets of 8 unique odor cues (sets A and B), which predicted reward based on one of two orthogonal rules (Fig. 1a). Other than the difference in rules predicting reward, which we will describe below, sessions were otherwise identical in the structure of the events in each trial and the distribution or potential rate of reward. Each trial began with a light signaling the rat to sample an odor at the designated port. When a nosepoke into the port was detected, one of the eight odors was delivered, requiring the rat to decide whether to respond to a nearby fluid well to obtain a reward (Fig. 1b). Responses on positive trials resulted in the delivery of 50 microliters of sucrose solution, followed by a 4-second light-off period before the initiation of a new trial. Withholding a response, regardless of trial type, resulted in no outcome and terminated the trial.

**Figure 1.**
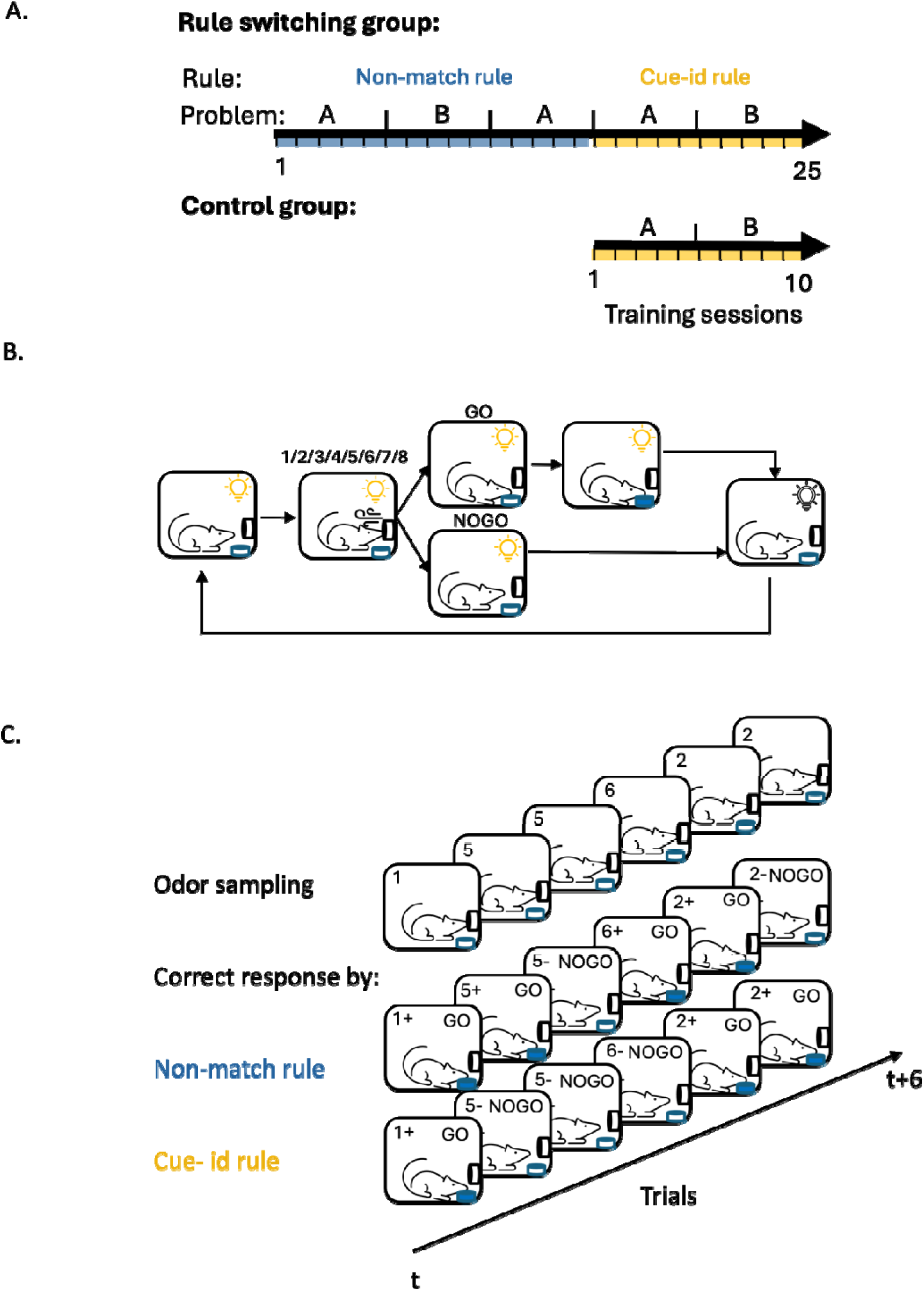
**A.** Curriculum overview. Rats were trained on a sequence of odor guided discrimination tasks governed by two orthogonal rules: ‘Non-match’ and ‘cue-id’. For each rule, two different odor sets, comprised of 8 different odors (problem A and B), were used for at least 5 consecutive sessions. Control group was trained solely on ‘cue-id’ rule. **B.** Schematic of trial structure. **C.** Schematic of the task design in 6 consecutive trials and the correct response according to the different rules. In each trial, one odor was presented (1-8) and the rat had to decide whether it is associated with reward.

During the initial training phase, the odor cues predicted reward based on a ‘non-match’ rule, where a reward was delivered if the response was to an odor different from the one sampled in the previous trial (Fig. 1c; “non-match rule”). In the first training session on this problem, the rats exhibited their default response of indiscriminately responding to the fluid well on all trials (Fig. 2a, ‘Non-match A 1^st^’). But then, in the following sessions, they gradually learned to respond correctly on ‘non-match’ trials and withhold their response on ‘match’ trials. After 9-10 sessions, they learned to respond only if the odor was different from the odor sampled in the previous trial (Fig. 2a ‘Non-match A last’). This response pattern resulted in increased behavioral accuracy based on the non-match rule (Fig. 2d, left; blue line) and a decrease in the number of trials required to reach 80% accuracy for successive sessions (Fig. 2e, left). After showing robust and stable performance (80% correct for 3 consecutive sessions), the rats were trained on a new problem where the rule remained the same, but eight new odors were introduced (Fig. 2a, ‘Non-match B). They successfully generalized the match/non-match rule to the new odors, reaching 80% performance within a single session (Fig. 2d-e right). Finally, the rats were retested on the original problem, demonstrating robust and persistent retention of prior learning (Supplementary fig. 1-Non-match A’).

**Figure 2.**
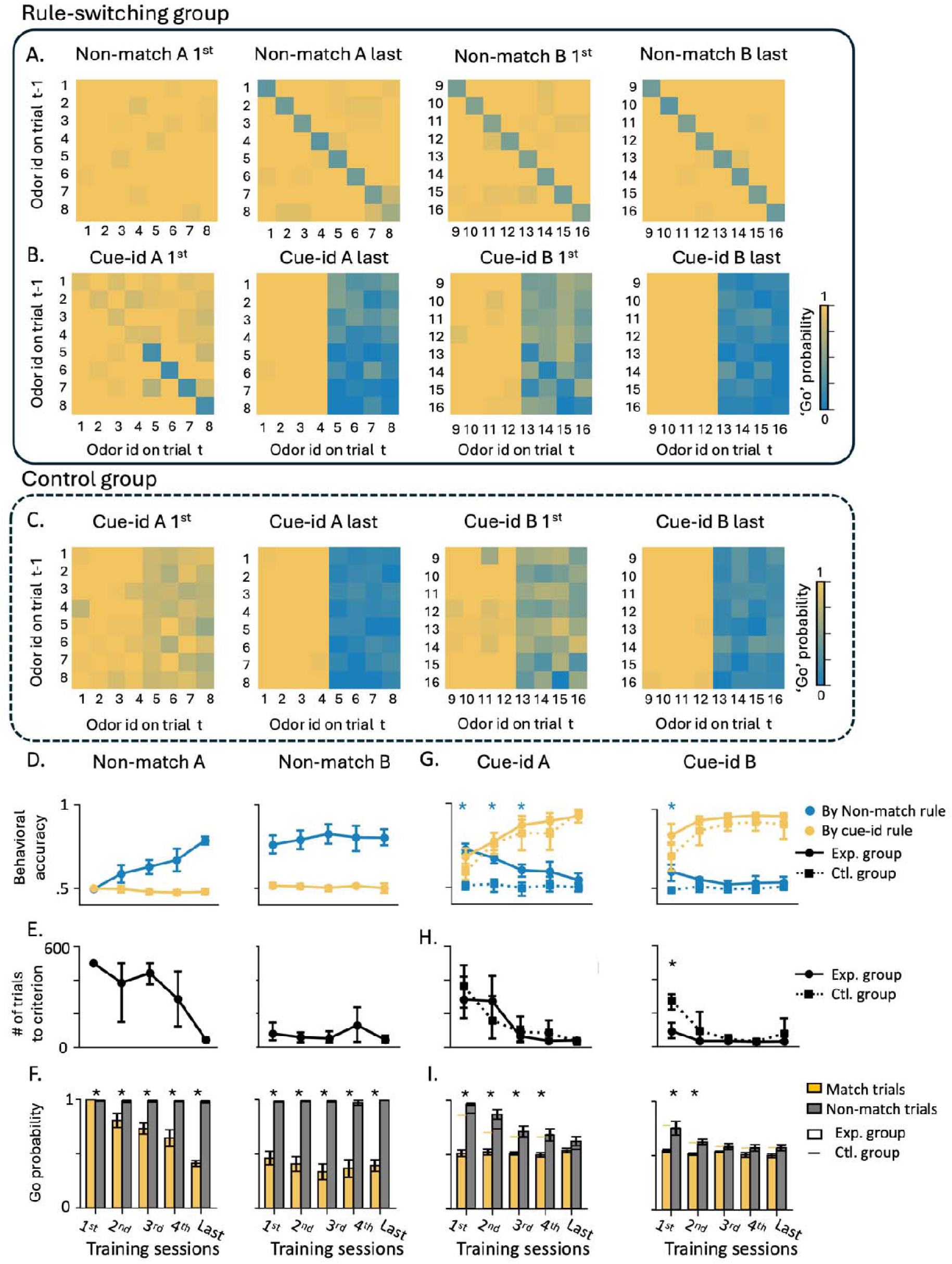
**A.** Probability of responding ‘go’ across all trial types (different odors in match/non-match configurations) in the first and last training sessions under the non-match rule for A and B problems. **B.** Same as A but for training under the cue-identity rule. **C.** Same as A-B, but for the control group, which was trained exclusively on the cue-identity rule. **D.** Accuracy for training under the non-match rule in A (left) and B (right) problems, according to the non-match rule (blue) and cue-identity rule (yellow) (mean ± SEM). **E.** Number of trials required to reach the learning criterion within each session under the non-match rule in A and B problems (see Methods for details). **F.** Go probability across different sessions under the non-match rule, separated into match (yellow) and non-match (gray) trials (mean ± SEM; * P < 0.05, Tukey’s Honest Significant Difference test). **G-H.** Same as panels D-E, but showing data from training sessions under the cue-identity rule for A and B problems. Solid lines represent the experimental group, while dashed lines represent the control group. Asterisks indicate statistically significant differences between groups (P < 0.05, Tukey’s Honest Significant Difference test). **I.** Same as panel F, but for training sessions under the cue-identity rule in A and B problems, comparing experimental (bars) and control (horizontal lines) groups. (* P < 0.05, Tukey’s Honest Significant Difference test).

After retesting on the original problem, rats began the second training phase, in which the odors remained the same, but the task rule changed to be based on ‘cue-identity’, where rewards were predicted by the identity of each odor rather than the match or non-match comparison with the prior trial (Fig. 1c; “cue-id rule”). Half of the odors (1-4) were associated with a potential reward (’rewarded odors’) while the other half (5-8) were not (‘non-rewarded odors’). The rats adapted their behavior to the new rule, learning to respond only to the rewarded odors and to withhold responses to the non-rewarded odors, regardless of the odor sampled in the previous trial (Fig. 2b, ‘Cue-id A’). This adaptation was gradual, resembling initial learning of the match/non-match rule (Fig. 2g left; yellow solid line, Fig. 2h left), however, unlike that initial learning, errors were not committed randomly. Instead, there was a higher probability of responding when the presented odor was a non-match to the prior trial (Fig. 2i left). This resulted in high residual accuracy according to the non-match rule (Fig. 2g left; blue solid line), particularly in the beginning of each session (Supplementary fig. 1). This pattern reemerged when the rats were presented with the next problem (Fig. 2b, ‘Cue-id B’); they again initially followed the old rule before fully committing to the new relevant rule (Fig. 2g-i right). Notably, this pattern was different from that in the initial phase of learning on the non-match rule, where the rats did not show any bias in their errors based on cue-identity (Fig. 2d; yellow lines). Thus, their bias to follow the now irrelevant orthogonal ‘non-match’ rule, after a new ‘cue-identity’ rule was introduced, depended on the prior experience. To confirm this dependency, we trained another group of rats exclusively on the ‘cue-identity’ rule (Fig. 1a “control group”). These rats also learned to follow the ‘cue-identity’ rule (Fig. 2c), gradually increased their behavioral accuracy according to this rule (Fig. 2g, yellow dashed lines) and reached the learning criterion with fewer trials (Fig. 2h, dashed line). Their behavior was not different in any aspect from the behavior of the rats that underwent the full learning curriculum, except that their errors were distributed randomly, rather than being more likely when the odor was a non-match to the prior trial (Fig. 2g, blue dashed lines). Overall, these results confirm that, when confronted with a new contradictory rule, the rats in our main experimental group showed residual effects of the prior behavioral schema, which gradually diminished as they learned to apply the new schema appropriate to the new rule.

### Single unit correlates during learning of orthogonal rules

We recorded single-units from the lateral orbitofrontal cortex (lOFC) during all of the training described above. The total number of neurons, number of neurons per rat, average firing rate, and percentage of responsive units remained stable throughout the training (Supplementary fig. 2). However, the selectivity of individual neurons to different task components changed according to the rule in effect. While most of the neurons recorded in the first training session had a significant response during the odor sampling period (Supplementary fig.2d, Fig.3a, ‘Non-match A, 1^st^ ‘), only a few neurons fired differently in rewarded versus non-rewarded trials. This is captured in Figure 3b, which shows the difference in the z-score versus baseline for rewarded versus non-rewarded trials for each unit (top) as well as the average z-score across all neurons (bottom). To quantify the proportion of neurons with significant selectivity to the rule, we used the parameter-free ZETA-test ^33^ to compare the neurons spike train in response to the odor sampling period in rewarded and non-rewarded trials (Fig. 3c). This analysis revealed that, in the first training session, less than 5% of the neurons were selective to the non-match rule. However, as learning progressed and the rule consolidated, more neurons had a different response in rewarded and non-rewarded trials (Fig. 3a-b, ‘Non-match A, last’), and the proportion of the non-match rule selective neurons gradually increased to 25% (Fig. 3c, ‘Non-match A’). This increased representation persisted as the schema generalized to a new problem with a different odor set (Fig. 3a-c; ‘Non-match B’).

**Figure 3.**
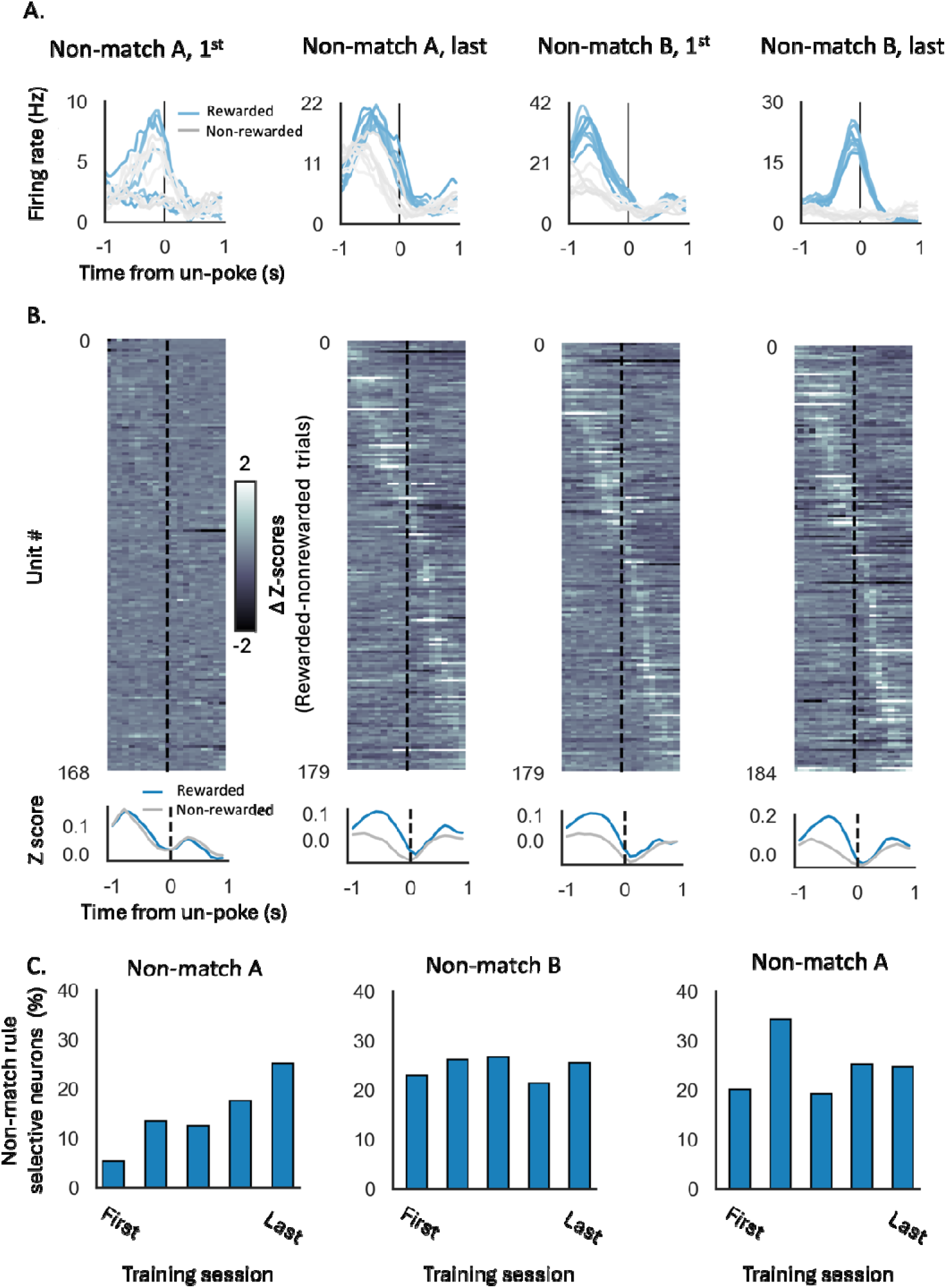
**A.** Peristimulus time histograms (PSTHs) of example neurons in response to the different odors in rewarded (blue) and non-rewarded (gray) trials. The PSTHs were aligned to the decision time (un-poke from odor port). **B. Top.** Difference in Z-score of all recorded units per session in rewarded versus non-rewarded trials, aligned to the decision time. **Bottom.** Z-score in rewarded (blue) and non-rewarded (gray) trials, averaged over all units. **C.** Proportions of neurons with a significant selectivity to the non-match rule, based on parameter-free ZETA-test (see methods).

To evaluate the degree of selectivity to the orthogonal ‘cue-identity’ rule during match/non-match sessions, we compared the difference in z-scores of individual units, as well as the proportion of selective neurons based on the ZETA-test comparison, dividing the trials to rewarded and non-rewarded trials according to both rules (i.e. either the current match/non-match or the future cue-identity rule). Selectivity to the future ‘cue-id rule’ was relatively low, declined in the first few sessions, and then remained low for the remainder of training on the ‘non-match rule’ (Fig 4a-b, ‘Non-match A’).

**Figure 4.**
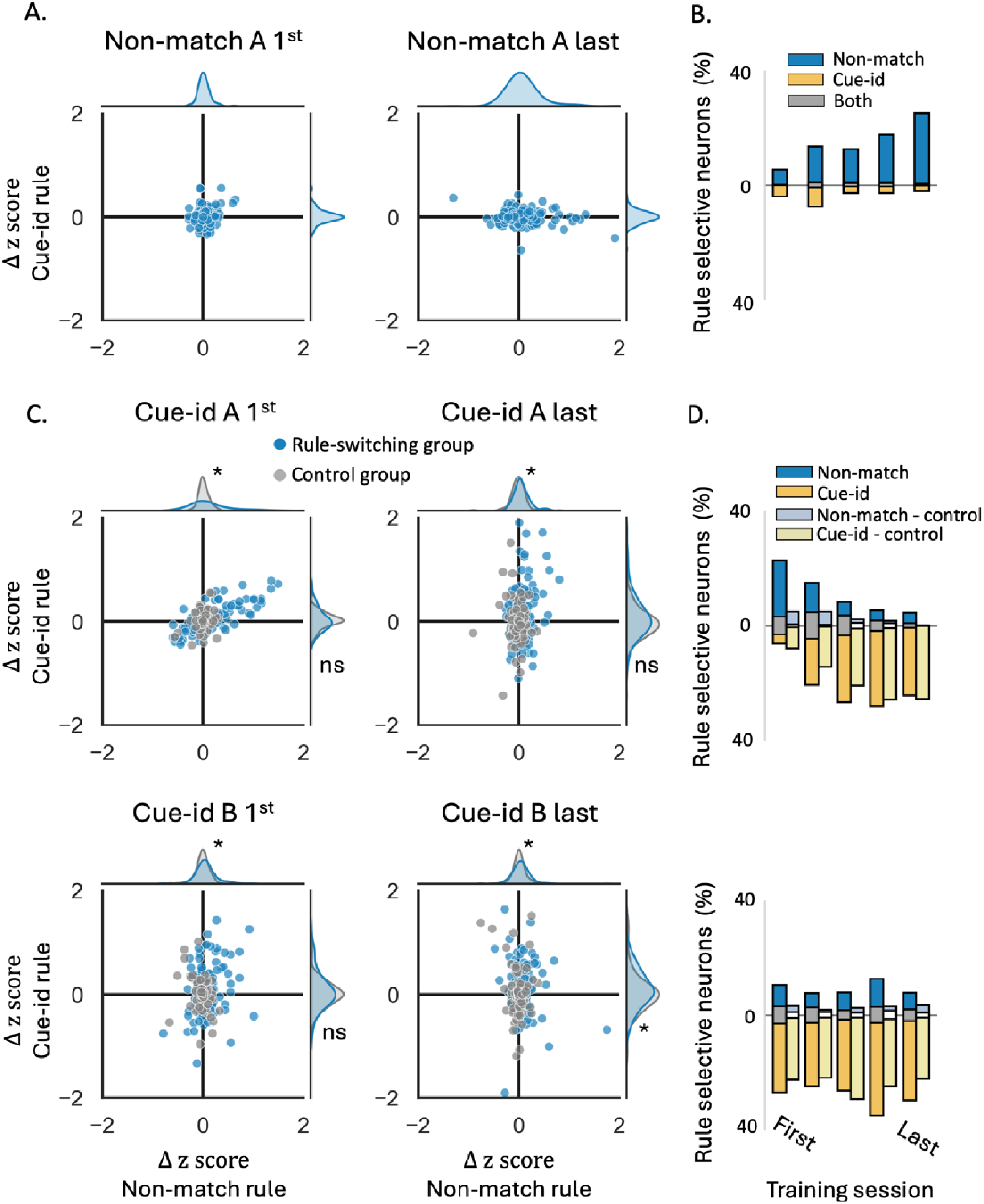
**A.** Delta z-score of individual units in rewarded vs. non-rewarded trials, plotted for the non-match rule (X-axis) and the cue-identity rule (Y-axis), during the first (left) and last (right) sessions of problem A under the non-match rule. Marginal axes display the distributions of delta z-scores for each rule. **B.** Proportions of neurons showing significant selectivity, for the non-match rule (blue), cue-identity rule (yellow), or both rules (gray) across different sessions of problem A under the non-match rule. **C.** Same as A, but for problems A (top) and B (bottom) under the cue-identity rule. Blue markers represent individual units from rats that underwent the full learning curriculum and first learned the non-match rule, while gray markers correspond to units from the control group, which was trained only on the cue-identity rule. Marginal axes show the distributions of delta z-scores for the rule-switching group (blue) and control group (gray). (* P < 0.05, Kolmogorov-Smirnov test comparing the two groups). **D.** Same as B, but for sessions under the cue-identity rule, comparing the rule-switching group (dark bars) and control group (light bars).

When cue-id training began, the representation gradually shifted to represent the new relevant rule (Fig. 4c-d, Supplementary fig.3). The difference in response to rewarded versus non-rewarded trials according to this rule was increased in many of the units (Fig. 4c, blue markers) and the proportion of neurons selective to this rule gradually increased (Fig. 4d, dark yellow bars). However, a significant proportion of neurons still exhibited a substantial difference in response to rewarded versus non-rewarded trials according to non-match rule (Fig. 4c) and remained selective to this irrelevant rule (Fig. 4d, dark blue and gray bars; 5%). This residual selectivity was observed even after behavior fully conformed to the new rule (Fig. 2g).

To confirm that this residual representation of the irrelevant rule was not due to intrinsic “mixed selectivity” or encoding of latent relationships, instead reflecting prior experience with the first rule, we compared it with the neural representation in the control group of rats that trained solely on the cue-id rule. In these rats, the difference in responses to rewarded versus non-rewarded trials according to the non-match rule was significantly smaller than those of the ‘rule-switching’ group (Fig. 4c, gray markers; p<0.01) with almost no units that were significantly selective to this irrelevant rule (Fig.4d, light blue and white bars), thus suggesting that prior behavioral schema implementation had a prolonged effect on the OFC representation.

### Patterns of population activity during learning orthogonal rules

To investigate the neural representation of conflicting behavioral schemas by the population activity in the OFC, we reduced the dimensionality of the neural responses using Uniform Manifold Approximation and Projection (UMAP), embedding the data into a three-dimensional space. The UMAP embeddings were plotted to provide a clear visualization of trial-specific neural representations, with data points distinguished by odor identity (color) and trial configuration (marker style). As rats learned to implement the first rule, the neural representation evolved to form distinct clusters of match and non-match trials based on the rule in effect (Fig. 5a left; ‘x’ and ‘o’ markers, respectively), rather than on the identity of the odor presented (different colors). To further quantify this separation, dendrograms constructed from the UMAP embeddings demonstrated strong clustering of trials by reward contingency under the ‘non-match’ rule (Fig. 5a left). When the same rule was generalized to the next problem, a similar population representation was observed, with trials continuing to show clear separation according to the non-match rule (Fig. 5a right). This consistency across problems highlights the stability of the neural representation when the task rule remains unchanged and the unimportance of cue identity in rats first trained on the non-match rule. As training progressed, the separation between odors in the match versus non-match configurations increased, indicating consolidation of the first behavioral schema (Fig. 5d, blue curves).

**Figure 5.**
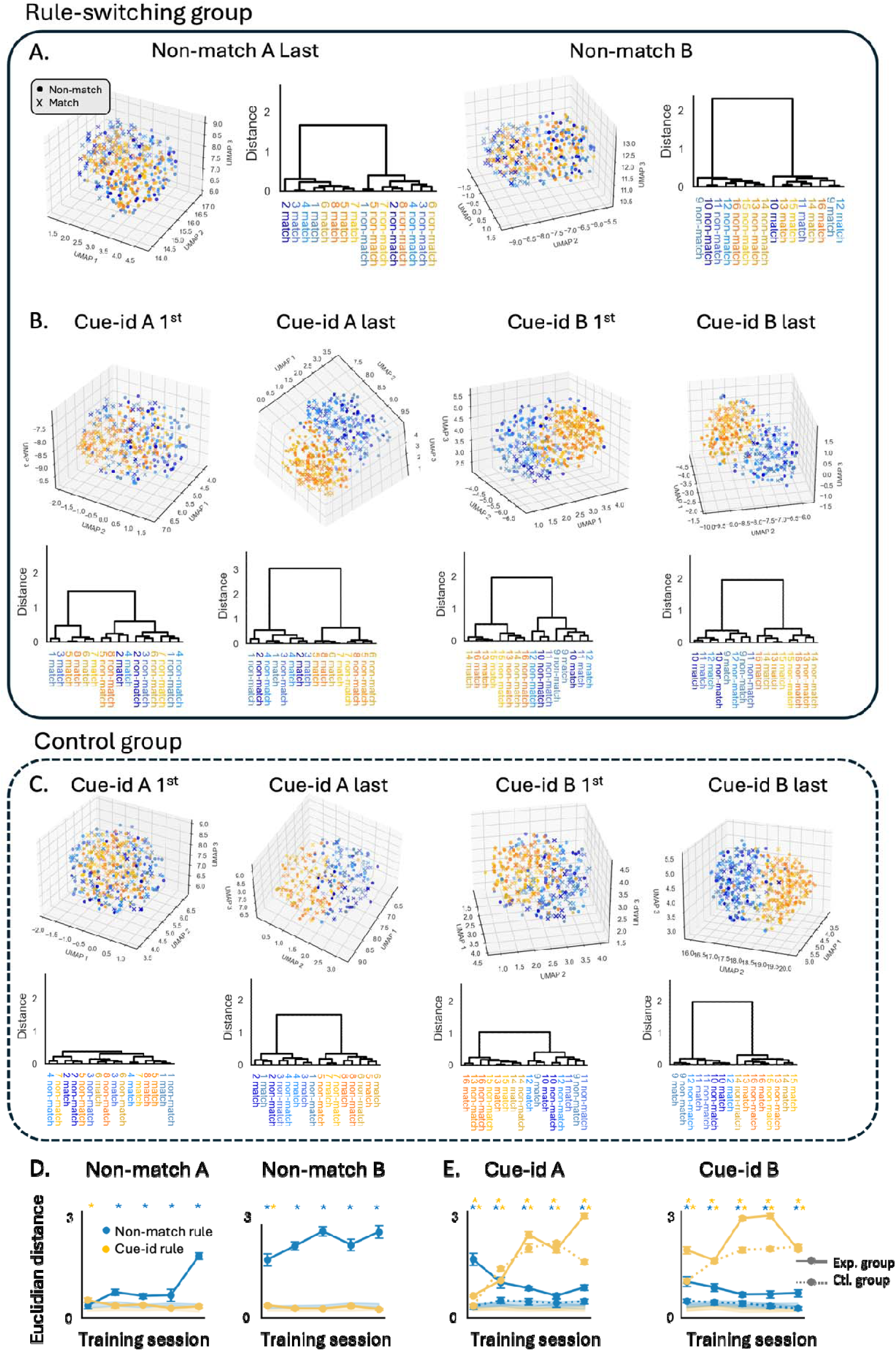
**A.** 3D UMAP visualization of neural population activity during the odor sampling period, with trials separated by odor identity (blue shades: odors 1–4, orange shades: odors 5–8) and trial configuration (’o’ for non-match trials, ‘x’ for match trials). Each raster represents pseudo-population responses from the final training session of A and B problems under the non-match rule. Adjacent dendrograms show trial-type clustering based on UMAP distances. **B.** Same as A, but for problems under the cue-identity rule. **C.** Same as B, but for the control group of rats. **D.** Euclidean distance in the UMAP space between pairs of odors in the match vs. non-match trial configurations (blue) and rewarded vs. non-rewarded odors (yellow) across different sessions under the non-match rule (mean ± SEM). Shaded error bars represent distances from a shuffled dataset. Asterisks indicate statistically significant differences from shuffled data for the non-match rule (blue) and cue-identity rule (orange) (p < 0.01, permutation test). **E.** Same as D, but comparing rats from the main rule-switching group (solid line) and the control group (dashed lines) across all sessions under the cue-identity rule. Asterisks in the upper row denote statistically significant differences for the control group.

When rats were confronted with the new ‘cue-id’ rule, the neural population activity in the OFC adapted to better separate trials according to this rule. In these sessions, the UMAP embeddings revealed a growing separation between rewarded and non-rewarded odors (Fig. 5b, blue and orange shades, respectively; Fig. 5e, solid orange curve), reflecting the implementation of the new behavioral schema. However, the neural representation also retained the previously learned ‘non-match’ rule, as evidenced by the partial separation of trials based on the irrelevant rule (Fig. 5b, ‘o’ and ‘x’ markers; Fig. 5e, solid blue curve). To confirm that the persistence of the old rule’s representation was not due to chance and reflected prior experience with the ‘non-match’ rule, we compared these findings to those in the control group trained exclusively on the ‘cue-id’ rule. In these rats, the UMAP analysis revealed a robust separation between rewarded and non-rewarded odors, similar to the separation observed in the experimental group (Fig. 5c). However, the representations of match and non-match trials remained overlapping, and were not different from a shuffled dataset (Fig. 5e, dashed blue curve).

The similarity in the patterns of activity in the neural activity space on different trial types can also be represented in a matrix-form, which can be used to understand how reliably information about different aspects of the task is represented, in this case the two rules. To illustrate this, we constructed three exemplar templates showing similarity based on each rule alone or in combination (Fig. 6a). Thus, in the ‘non-match template’, similarity is high for odors presented in non-match configuration or for odors presented in the match configuration, whereas in the ‘cue-id template’, similarity is high for rewarded odors or non-rewarded odors, regardless of whether they were presented in match or non-match configuration. Finally, in the ‘both-rules template’, the similarity of the activity pattern is high between rewarded odors or between non-rewarded odors, but only if they also share the same match/non-match trial configuration. We compared those templates to the results from an analysis of neural population firing rates during the odor sampling time during learning across sessions involving the two rules. To quantify the response similarity between different trial types, we employed a Support Vector Machine (SVM) decoder, trained to predict the trial type based on a vector of firing rates for the neurons in the population,for each rat and session separately. We utilized a leave-one-out cross-validation strategy to assess the accuracy of the decoders and plotted its predictions as confusion matrices (Fig. 6b-d). As rats learned to implement the first rule, the decoders increasingly confused trial types that shared the same potential outcome (match/non-match; Fig. 6b) and became similar to the ‘non-match template’ (Fig. 6e, blue lines, see methods). In contrast, the similarity to the ‘cue-id template’ or to the ‘both-rules template’, remained low (Fig. 6e, yellow and gray lines).

**Figure 6.**
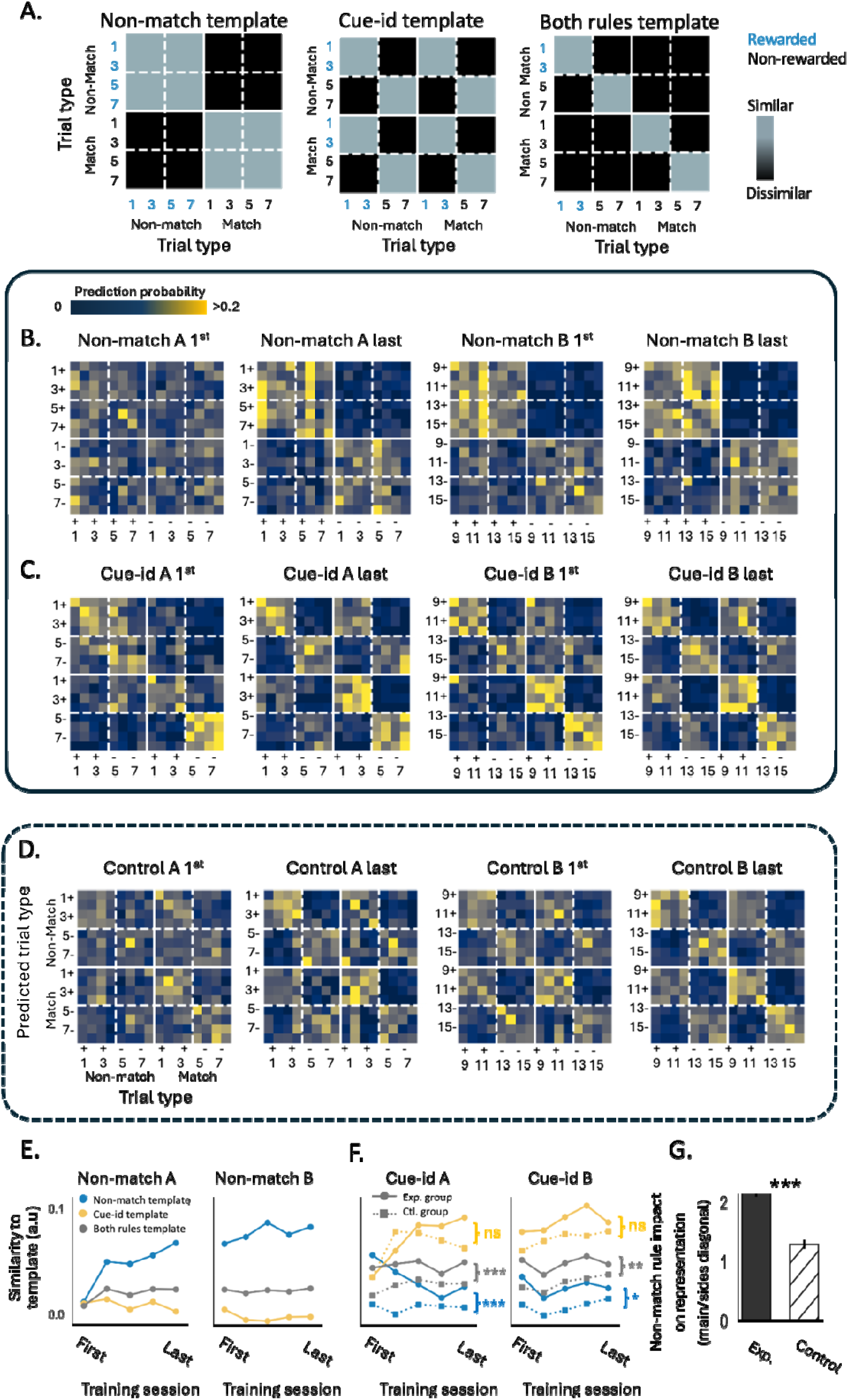
**A.** Hypothesized templates of confusion matrices, illustrating expected patterns of similarity between rewarded trials, based on reward distinction according to the non-match rule, cue-identity rule, or both rules. **B.** Confusion matrices, averaged across rats, showing the probability of a support vector machine (SVM) decoder predicting the trial type based on neural population activity during the first and last training sessions of A and B problems under the non-match rule. Each matrix row shows the probability of predicting each of the possible trial types (columns), given the neural population activity for a given trial type (row). **C.** Same as B, but for problems under the cue-identity rule. **D.** Same as C, but for the control group of rats. **E.** Similarity of population decoding confusion matrices across different rats in all sessions under the non-match rule, compared to the ‘Non-match template’ (blue), ‘Cue-identity template’ (yellow), and ‘both rules template’ (gray). **F.** Same as E, but comparing rats from the main group (circles) and the control group (squares, dashed) in all sessions under the cue-identity rule. **G.** Impact of the non-match rule on population representation during cue-identity sessions, calculated as the ratio between probabilities in the main diagonal and side diagonals of the confusion matrices (*** p < 0.001, Wilcoxon test).

When confronted with the new rule, the pattern of neural activity began to reflect the outcome according to the new rule (Fig. 6c) and became better aligned with the template corresponding to the ‘cue-id’ rule (Fig. 6f, solid yellow lines). However, the similarity between trials that predicted reward by both rules was higher, meaning it more likely for the decoders to be confused within these trial types (Fig. 6c). Consequently, while the similarity to the ‘cue-id template’ increased, the similarity to the ‘non-match template’, which was based on the former rule, or to the ‘both-rules template’ reflecting the integration of the two rules, remained significantly above chance over many sessions involving both problem A and problem B (Fig. 6f, solid blue and gray lines).

This result contrasts with the findings in the control group of rats, where the population activity in OFC converged, with training, to become better aligned with the ‘cue-id’ template (Fig. 6d, Fig. 6f; dashed yellow lines) with little or no influence of the irrelevant match-non-match rule (Fig. 6f; dashed blue and gray lines). This difference is most striking in a direct comparison of classification along the main and the side diagonals of the confusion matrices after cue-identity training in both groups, which were similar in controls but asymmetric in the experimental group (Fig. 6g). Analysis of neural decoders that trained to predict reward according to each rule further confirmed that the residual representation of the irrelevant rule was unique to the group of rats who previously learned that rule (Supplementary fig. 4a) and was not an artifact of the animal selection (Supplementary fig. 4b).

Overall, these findings, derived using multiple approaches, converged on the same conclusion. First, both individual neurons in the OFC and the pattern of activity across ensembles of OFC neurons dynamically adapt to encode the relevant behavioral schema. Second, they do so while retaining significant traces of the previously acquired rule, even long after behavior had fully conformed to the new rule across multiple problems.

### Effects of residual encoding on optimal behavior during learning of orthogonal rules

Interleaving old information with new can allow the old information to remain available to be used in future scenarios, however this practice is generally assumed to have contrary effects on behavior. For example, the representation of previously acquired rules would be expected to interfere with a successful acquisition and expression of a new conflicting rule. If this is true, we would expect to find a positive correlation between encoding of the previous rule and the tendency of the rat to follow that irrelevant rule. Alternatively, it is also possible that the network is able to represent the previous rule in parallel with the new learned rule, without any direct effect on the behavior or the acquisition of the new rule. Or, finally, it is possible that representation of the previous rule is beneficial to acquisition of the new. Indeed, there are even some examples, such as the over-training reversal effect ^34, 35^, which suggest that strong representation of a prior contradictory rule can facilitate new learning.

To adjudicate between these possibilities, for each rat and each session, we computed the correlation between 1) the accuracy of the neural decoder at distinguishing between potential outcomes based on the two rules, and 2) h the accuracy of the behavior according to each rule. In the first learning phase, when only exposed to one rule, behavioral accuracy was significantly correlated with the decoder accuracy based on the relevant non-match rule (Fig 7a, blue markers). The better the neural decoder distinguished between match and non-match trials, the more accurately the rat followed that rule and correctly responded only to non-match trials. During this training phase, the correlation between the tendency to perform based on the cue-id rule, and how well this rule has been represented in the neural activity, was low or non-significant (Fig 7a, yellow markers).

**Figure 7.**
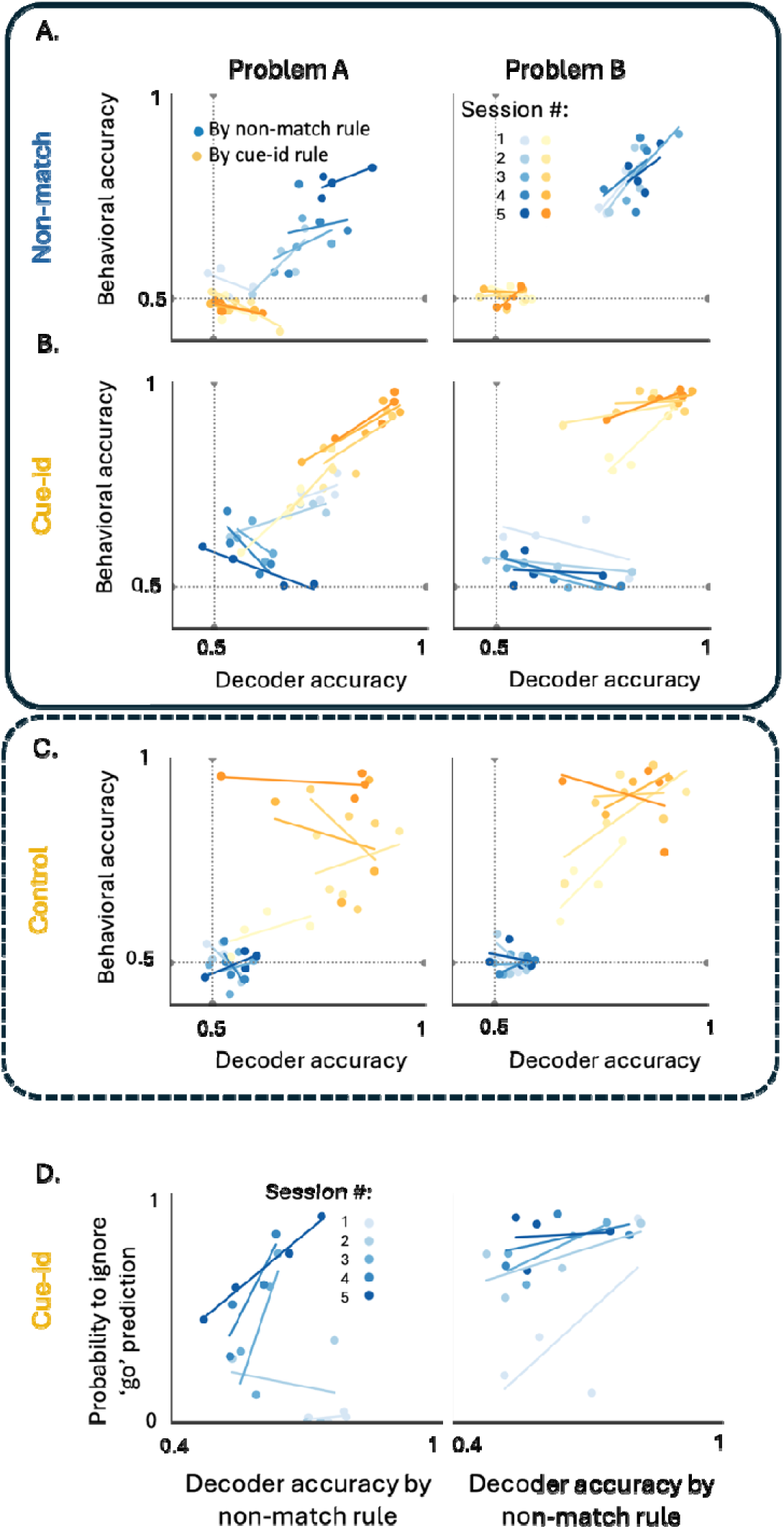
**A.** Correlation between behavioral accuracy and population decoder accuracy according to the non-match rule (blue) and the cue-id rule (yellow), in problem A (left) and problem B (right) under the non-match rule. Shades represent different sessions in each problem. **B.** Same as A but for problems A (left) and B (right) under the cue-id rule. **C.** same as a-b but for control group who trained solely on cue-id rule. **D.** Correlation between the decoder accuracy of the irrelevant rule and the proportion of trials that were labeled as go trials by this decoder but have been ignored by the animal who chose to hold their response, as expected by the new relevant rule.

When confronted with the new cue-identity rule in the second phase of training, the first two sessions were characterized by a positive correlation between the accuracy of the neural decoder and of the behavioral response, for each of the two rules. As learning progressed and the two rules were distinguished by behavior (Fig 2), the correlation between the neural decoding and the behavioral performances of the new relevant rule remained positive (Fig 7b, yellow markers), but the correlation between these measures inverted for the irrelevant non-match rule. That is, the better the decoder distinguished between rewarded and non-rewarded trials according to the non-match rule, the less the rats followed this rule in their responses (Fig. 7b, blue markers) and the more they followed the new rule. This inverse relationship was not observed in the control group of rats who were not trained on the non-match rule (Fig.7c).

To directly investigate the relationship between the neural representation of the irrelevant rule and the behavioral response, on a trial-by-trial basis, we focused on the incongruent trials where reward is expected according to the non-match rule but not according to the new rule. For these trials, we calculated the proportion of trials that were correctly labeled as rewarded trials by the non-match neural decoder, but where the rats nevertheless chose not to respond, in accordance with the new relevant cue-identity rule. This probability of ignoring the ‘go’ prediction of the old rule increased as training on the new rule progressed, and, surprisingly, was positively correlated with the fidelity of the neural representation of the old rule (Fig 7d). In other words, the more robust and faithful the representation of the old irrelevant rule, the more expert the rat became at ignoring its prediction, instead acting in agreement with the prediction of the new relevant rule. These results indicate not only the absence of any negative effect of interleaving the prior rule on optimal choice behavior, but actually show a facilitative effect.

## Discussion

Schemas are critical to efficient behavior in the world but can also introduce problems when they become irrelevant to new environmental conditions ^19, 36^. Growing evidence points to a key role for the OFC in forming and deploying schemas in new situations congruent with previously acquired knowledge ^19, 20, 37, 38^. But how does this role affect the learning of a new and conflicting behavioral schema? Is representation of the prior schema in the OFC a source of interference, slowing or disrupting learning until it is erased, or do representations in the OFC coexist or even facilitate learning of new contradictory information? Here we found evidence consistent with the latter proposal. When rats were asked to acquire a second schema that conflicted with previous learning, OFC neurons interleaved the new information with the old. This mixed representation was not observed in control rats trained only on the second schema, and persisted even after the rats had shown expertise on new problems of the second type. Further, when encoding was examined during initial learning of the second, conflicting schema, clarity of representation of the prior schema was correlated with correct performance on the new one. That is, the more strongly OFC neurons represented the prior schema, interleaved with the new, the better the rats performed on the new schema. This indicates not only that OFC is not a source of interference when conflicting schemas must be resolved, but further that it is functioning to support this resolution by accurately and independently representing the old rule as the new is being acquired.

Recent studies have provided insights into the neural mechanisms underlying schema formation and implementation, revealing a dynamic interaction between the hippocampus and prefrontal, especially orbitofrontal, areas in assimilating new experiences into preexisting networks of associations ^15, 16, 39–46^. Neural ensembles in these regions appear to converge into a hierarchical organization, structuring relationships between overlapping elements within the task space ^20, 38, 47–50^. The assimilation of information that aligns with prior knowledge is accelerated by reactivating neuronal ensembles that represent the relevant schema, adjusting activity to incorporate novel details while preserving the low-dimensional structure relevant to common task demands ^43, 44, 51, 52^. Our findings further highlight the key role of the OFC in this process. Individual neurons in the OFC and the population activity patterns across OFC ensembles dynamically adapted to quickly encode the common features of new problems of a type. Specifically, neuronal responses became more similar for stimuli associated with the same outcome, reflecting the OFC’s role in organizing task-relevant information. However, as noted above, our results also show that the persistent representation of a prior schema in the OFC facilitates the detection of shifts in task demands that necessitate the formation of a new schema.

This role for the OFC in facilitating the learning of conflicting information is reminiscent of the long-recognized importance of the OFC to so-called cognitive flexibility ^53, 54^, epitomized by reversal learning. The OFC has been found to be necessary for reversal learning in many, though not all, settings ^24, 55–61^. The current results provide a more specific basis for this involvement in the accurate representation of the prior rule, independent of the new, so that errors or mistakes can be effectively signaled even while the new rule is acquired. Such a role should be particularly important when changes occur against a background of strong priors, since it is under these conditions, epitomized by the over-training reversal effect ^34, 35^, that representation of the prior information would be most effective. Accordingly, the OFC is most critical for initial reversal learning, and for reversal learning when contingencies have been stable for long periods ^55, 56^. When contingencies are rapidly changing, the OFC is unnecessary for accurately tracking the best choice, and it has even been shown to hinder reversal under these conditions ^56, 60, 62, 63^.

The current results provide a novel explanation for the involvement of the OFC in cognitive flexibility under these conditions in the persistent multiplexing of the schemas - the generalized cognitive maps ^64^ - deployed to guide behavior. This multiplexing is superior to forgetting in several ways. Most obviously, multiplexing ensures prior knowledge remains accessible for future transfer, allowing flexible adaptation if new problems fitting the old schema are encountered ^65^. However, in addition to this valuable function, multiplexing the old schema independently from the new also makes it possible for the old information to support strong error signaling to drive learning, even as the new information is laid down, whereas if the two rules were confused or the old was eliminated, teaching signals would weaken more quickly. In this light, it is interesting to consider that the OFC is not generally necessary for established performance. OFC manipulations have been shown repeatedly not to impact task performance after the relevant relationships and rules have been learnt; this includes settings such as economic choice, in which core functions of the OFC are purported to be at risk ^66, 67^. Instead, the OFC appears most necessary during learning or when new information must be acquired and used to change behavior ^68–75^. This suggests that the OFC is typically functioning as a follower in using existing information, and its role becomes decisive when integrating new and conflicting information which is required for normal behavior. Viewed from this perspective, the ability to hold two independent sets of information online for comparison and for learning becomes a core function of the OFC. Consistent with this, the OFC has broad influences on downstream areas, both supporting the representation of associative information in subcortical regions like amygdala and striatum ^76–82^, while also contributing to error signaling by midbrain dopamine neurons ^83–88^. If the multiplexed information evident here in OFC ensembles can be demixed through selective projections or downstream filtering, the OFC would be in a powerful position to simultaneously serve both roles ^89, 90^.

Notably, in this regard, the operation of OFC contrasts with that of standard artificial learning systems, which frequently experience catastrophic forgetting, where newly acquired information disrupts or erases previously stored knowledge, leading to a marked decline in performance ^91^. Although several strategies have been proposed to address this issue, they typically depend on computationally intensive architectures that lack the energy efficiency of biological neural systems. For example, ‘AlphaGo Zero’, an artificial neural network that achieved superhuman performance in the game of Go, still operates at a power consumption two orders of magnitude higher than the human brain’s modest 20-watt power budget. ^92–94^. Gaining deeper insights into the neural computations that facilitate seamless knowledge transfer in the brain could inspire more efficient and adaptable artificial intelligence, bridging the gap between biological and artificial learning ^95^.

## Methods

### Experimental Model and Subjects

This study used eight male Long-Evans rats (Charles River), weighing between 300 and 400 g and approximately 4 months old. The rats were housed individually under a 12-hour light/dark cycle at the AAALAC-accredited animal facility of the National Institute on Drug Abuse Intramural Research Program (NIDA-IRP), with unrestricted access to food. Water was removed the day before testing sessions, and the rats were allowed 10 minutes of water access in their home cages after each testing session. On days without testing, the rats had free access to water. All procedures were carried out in compliance with the guidelines of the U.S. National Institutes of Health (NIH) and were approved by the Animal Care and Use Committee (ACUC) of NIDA-IRP.

### Stereotaxic Electrode Implantation

Rats were implanted with 2-4 drivable electrode bundles, each containing 16 nickel-chromium wires (25 μm diameter, AM Systems, WA), totaling 32-64 electrodes, targeting the lateral orbitofrontal cortex (lOFC) at coordinates AP: 3 mm and ML: 3.2 mm. The electrode bundles were embedded in 27-gauge stainless-steel hypodermic tubing and mounted in a custom-built, 3D-printed microdrive. Before surgery, the bundles were trimmed to 1-2 mm with fine bone-cutting scissors (Fine Science Tools, CA) and spaced to maintain at least 25 μm separation between wires. During surgery, rats were anesthetized with isoflurane (3% induction, 1%–2% maintenance in 2 L/min O2) and secured in a stereotaxic frame (Kopf Instruments, Tujunga, CA) for electrode implantation. Electrode tips were initially positioned 4.2 mm ventral to the brain surface. Headcaps were affixed using 0-80 1/8” machine screws and dental acrylic, then encased in a custom 3D-printed protective cover. Post-surgery, rats received Cephalexin (15 mg/kg) orally twice daily for two weeks to prevent infection.

### Odor-Guided Discrimination Tasks

Behavioral training was conducted in aluminum boxes (∼18 inches per side) equipped with an odor delivery port and a sucrose solution well. Task execution was controlled by a custom C++ program controlling relays and solenoid valves, with infrared sensors detecting entries into the odor and fluid ports. Each trial began when two house lights were illuminated above the odor port, prompting the rat to nosepoke within 5 seconds. Upon entry, a 500 ms delay was followed by odor presentation, during which the rat was required to remain in the port for an additional 300 ms; early withdrawal aborted the trial. After this period, rats could exit the port, stopping odor delivery, and had 2 seconds to make a response at the fluid well. For rewarded trials, a response triggered a 50 μL sucrose solution (5% w/v) delivery after a 1000 ms delay. If no response occurred or the trial was non-rewarded, the house lights turned off, initiating a 4-second inter-trial interval.

Before odor training, rats were shaped to nosepoke and respond at the well for a reward. After 3-4 shaping sessions, they were trained on a series of odor-guided discrimination problems involving two sets of 8 unique odors (A and B), each set predicting reward based on one of two orthogonal rules (Fig. 1a). The first rule was a non-match rule, under which a response was rewarded if the odor presented on the current trial differed from that presented on the previous trial. Rats trained for at least 5 sessions on each problem before advancing to a new problem with the same rule but different odors. In the second phase, the task rule changed to a cue-id rule, under which the match comparison became irrelevant and rewards were contingent only on the identity of the odor presented on the current trial. Here, half of the odors (1-4/9-12) were associated with a potential reward (’go’ odors), while the remaining odors (5-8/13-16) were not (’no-go’ odors).

### Single-Unit Electrophysiology

Neural recordings were obtained using the Plexon OmniPlex system (v2.7.0; Plexon, Dallas, TX). Neural signals were digitized, amplified, and bandpass filtered (250 – 8,000Hz) to isolate spike activity. Thresholds were manually set on each channel to capture unsorted spikes. Behavioral timestamps were synchronized with neural data in real time. After recording, spikes were sorted offline using Offline Sorter (v4.0; Plexon, Dallas, TX). Single units were isolated in 2D feature space (PC1, PC2, nonlinear energy), after which unit and event timestamps were exported to Matlab for further analysis. The sorted data were exported for analysis in MATLAB (2021a; MathWorks). Electrodes were advanced approximately 120 µm between odor discrimination tasks to sample new neuronal populations, although neuron identity across sessions was not assumed.

### Behavior analysis

Behavioral data were collected using custom software written in C++, which sent event timestamps to the electrophysiological recording system. Raw data were processed in MATLAB 2021a (Mathworks, Natick, MA) to extract the time spent in the odor and reward ports relative to trial initiation. Further analyses performed using Python 3.9 and Jupiter notebook. Behavioral accuracy was quantified by the percent of trials on which the rats responded correctly according to the non-match rule or the cue-id rule. Number of trials to reach learning criterion was calculated as the number of trials in each session it took to cross 80% correct across 30 trials. Group differences were assessed with statsmodel library by ANOVA, with significant results followed by Tukey’s Honest Significant Difference post hoc test for pairwise comparisons at an alpha level of 0.05.

### Single unit analysis

The spike train for each isolated single unit was aligned to the decision time (time of un-poking from the odor-port). Spike number was counted with a bin of 50 ms. A peri-stimulus time histogram (PSTH) was generated by calculating the mean neural response across different trials. For each trial type, the mean response was smoothed using a convolution of a moving average filter, defined as a uniform filter with a window size of 5 time bins.

For each single unit, z-scores were calculated separately for rewarded and non-rewarded trials by normalizing the mean firing rate in the response window (500 ms prior to odor un-poke) against baseline activity (1 second prior to trial initiation). The z-score for non-rewarded trials was then subtracted from that of rewarded trials to obtain the delta z-score, representing the differential standardized activity between trial potential outcomes. To assess rule-specific neural modulation, each single unit’s delta z-score was computed separately for trials based on the two underlying rules.

To compare rule-dependent selectivity across the population, we applied the parameter-free ZETA test to identify neurons with significantly different firing rates in the window between poke and un-poke from the odor port for rewarded versus non-rewarded trials^33^. Briefly, the neuronal activity in response to the odor sampling period was compared against a shuffled null distribution created by permuting the activity across trials. This algorithm identifies responsive time windows without predefined parameters, ensuring that only statistically significant deviations from the null model are classified as true responses. Neurons that exhibited a significant evoked response and differential firing across these two conditions were labeled as selective for that rule. This process was conducted separately for the non-match and cue-id rules.

### Population analysis

To explore and quantify the representation of the learned rules by the neural population activity, we employed Uniform Manifold Approximation and Projection (UMAP), a non-linear dimensionality reduction technique, to embed neural activity into a low-dimensional space^96^. Neural firing rates were extracted from the 500 ms period preceding the un-poking from the odor port and normalized using a standard scaling method to ensure consistency across datasets. Recordings from equivalent training sessions of different rats were normalized independently, and the data were subsequently aligned by trial type to generate pseudo-ensembles. UMAP, implemented using the Python library umap-learn, was then applied to the aligned dataset, resulting in three-dimensional embeddings. These embeddings were visualized for each trial, with data points distinguished by odor identity (color) and trial configuration (marker style). To further quantify the relationships between the neural representation of different trial types in the UMAP space, we calculated the distances between their centroids and visualized these relationships using a dendrogram. For each trial type, the mean embedding was computed by averaging the UMAP dimensions for all data points within that trial type. The distances between these centroids were then calculated using a normalized Euclidean distance metric, which accounted for variance within each trial type. Hierarchical clustering was performed using the Ward linkage method, implemented in the Python library scipy, to generate a linkage matrix, which was subsequently used to construct a dendrogram. This dendrogram provided a hierarchical visualization of the relationships and separability of task-specific neural activity patterns in the UMAP space.

Additionally, the distances between the representations of odors in the match versus non-match configurations and the distances between odors across the ‘rewarded odors’ and the ‘non-rewarded odors’ groups were calculated for each session. To assess the significance of these distances against a null hypothesis of no difference between the two groups of odors, we performed a permutation test by shuffling trial labels within each session and recalculating the distances for the shuffled data. The observed distances were compared against this permutation null distribution of shuffled distances to determine whether the separation between rewarded and non-rewarded trials according to each rule was statistically significant for each session.

To assess the alignment of the neural population activity with each rule, we employed a support vector machine (SVM) classifier from scikit-learn library. The SVM model was trained separately for each animal and training session to predict trial type based on neural responses during the odor sampling period. A leave-one-out cross-validation strategy was applied to generate trial-wise prediction probabilities, which were visualized through confusion matrices. These matrices where then compared against the template matrices to calculate similarity to each template. To test the significance of similarity between the neural confusion matrix and rule-based templates, against a null hypothesis of the similarity to each template happening by chance, we conducted a permutation test (1,000 permutations). For each permutation, we randomly shuffled rows and columns of the neural confusion matrix to create a permuted matrix, then computed its similarity score with each target template by applying a scoring matrix. After calculating actual similarity scores for the unshuffled matrix, we compared these to the permutation-based scores to obtain p-values, indicating the proportion of permuted scores meeting or exceeding the observed score. This non-parametric test evaluated the alignment between neural activity and each rule template.

To further assess the decodability of each rule from the neural population, we trained additional classifiers to differentiate between rewarded and non-rewarded trials based on each rule. Using 1,000 permutations, we performed a permutation test to evaluate classifier accuracy as the mean of cross-validated scores and calculated an empirical p-value for statistical significance.

To isolate rule representation from the potential influence of animal choice, we regressed out choice-related variance from the neural data. This was achieved by using a linear regression model, where animal choice (treated as a predictor variable) was fitted to the neural response data. The residuals, representing neural activity after accounting for choice, were then used for subsequent SVM decoding analysis. This approach ensured that the classification accuracy reflected rule-specific neural patterns independent of the animal’s choice behavior.

The probability of ignoring the ‘go’ prediction was calculated as the proportion of trials that were labeled as rewarded trials by the ‘non-match’ classifier, but had resulted not resulted in a ‘go’ action.

## Acknowledgements

This work was supported by the Intramural Research Programs at the National Institute on Drug Abuse and the National Institute of Mental Health. The opinions expressed in this article are the authors’ own and do not reflect the view of the NIH/DHHS. The authors have no conflicts of interest to report.

## Data and code availability

The data described in this study and custom analysis code available on GitHub at https://github.com/IdoMaor/Persistent-representation-of-a-prior-schema-

---

**Supplementary 1.**
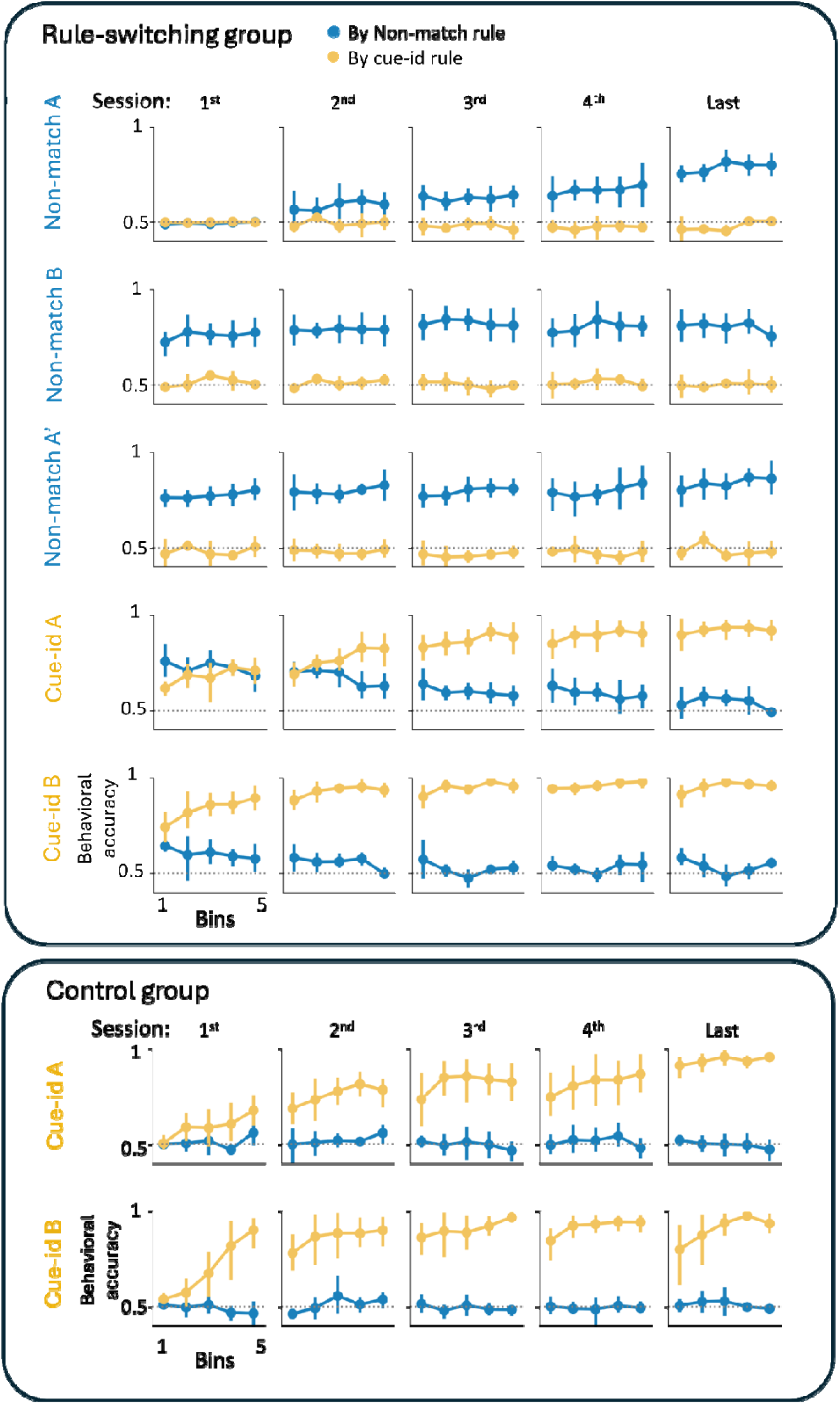
Accuracy according to the non-match (blue) and cue-identity (yellow) rules broken down to different bins within a session.

**Supplementary 2.**
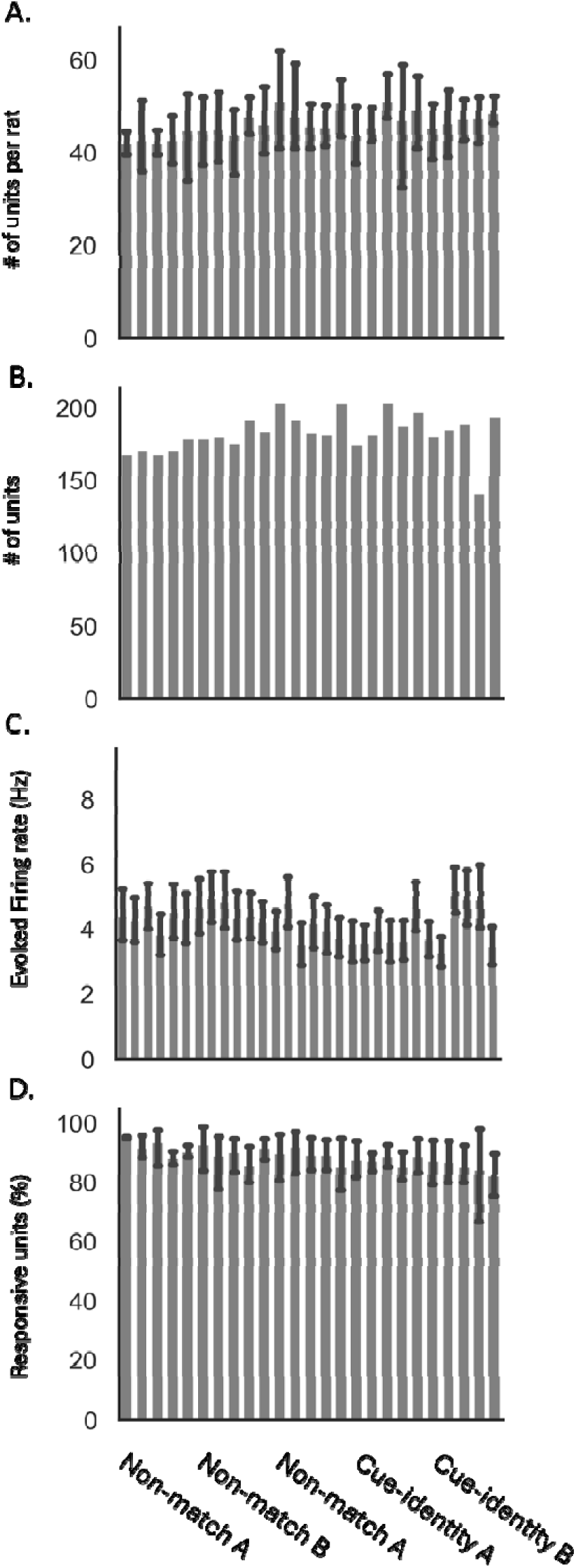
**A.** single units recorded per rat per session (mean+sem). **B.** Total number of single units recorded per session. **C.** Firing rate in 500 ms preceding the decision per session (mean+sem). **D.** Percentages of units with a significant response during the odor sampling period (mean+sem).

**Supplementary 3.**
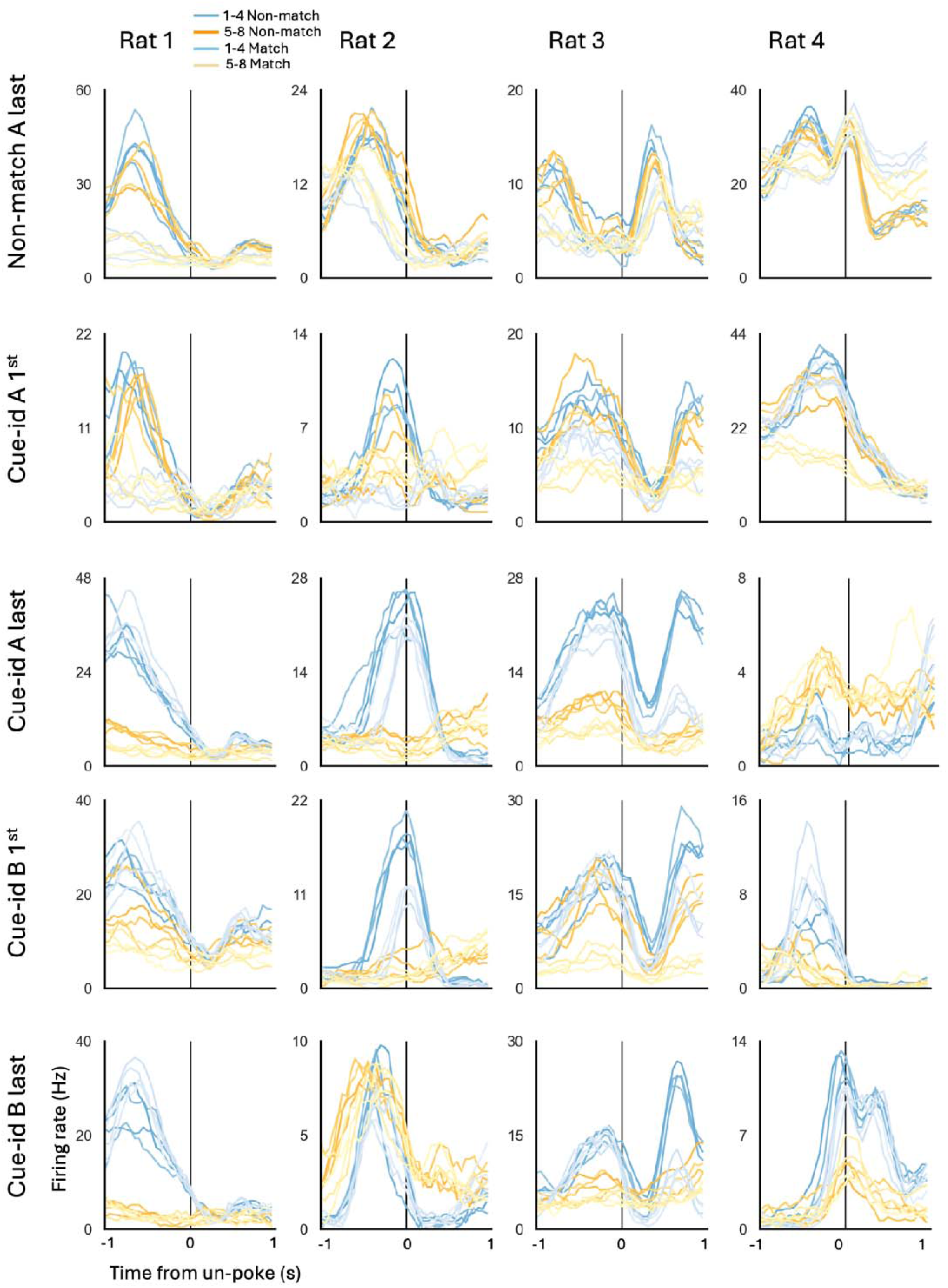
Peristimulus time histograms (PSTHs) of example neurons, from 4 rats and 5 learning sessions, in response to 16 different trial types: 8 odors (1-4: blue lines; 5-8: yellow lines) X 2 trial configurations (match: light colors; non-match: dark colors. The PSTHs were aligned to the decision time (un-poke from odor port).

**Supplementary 4.**
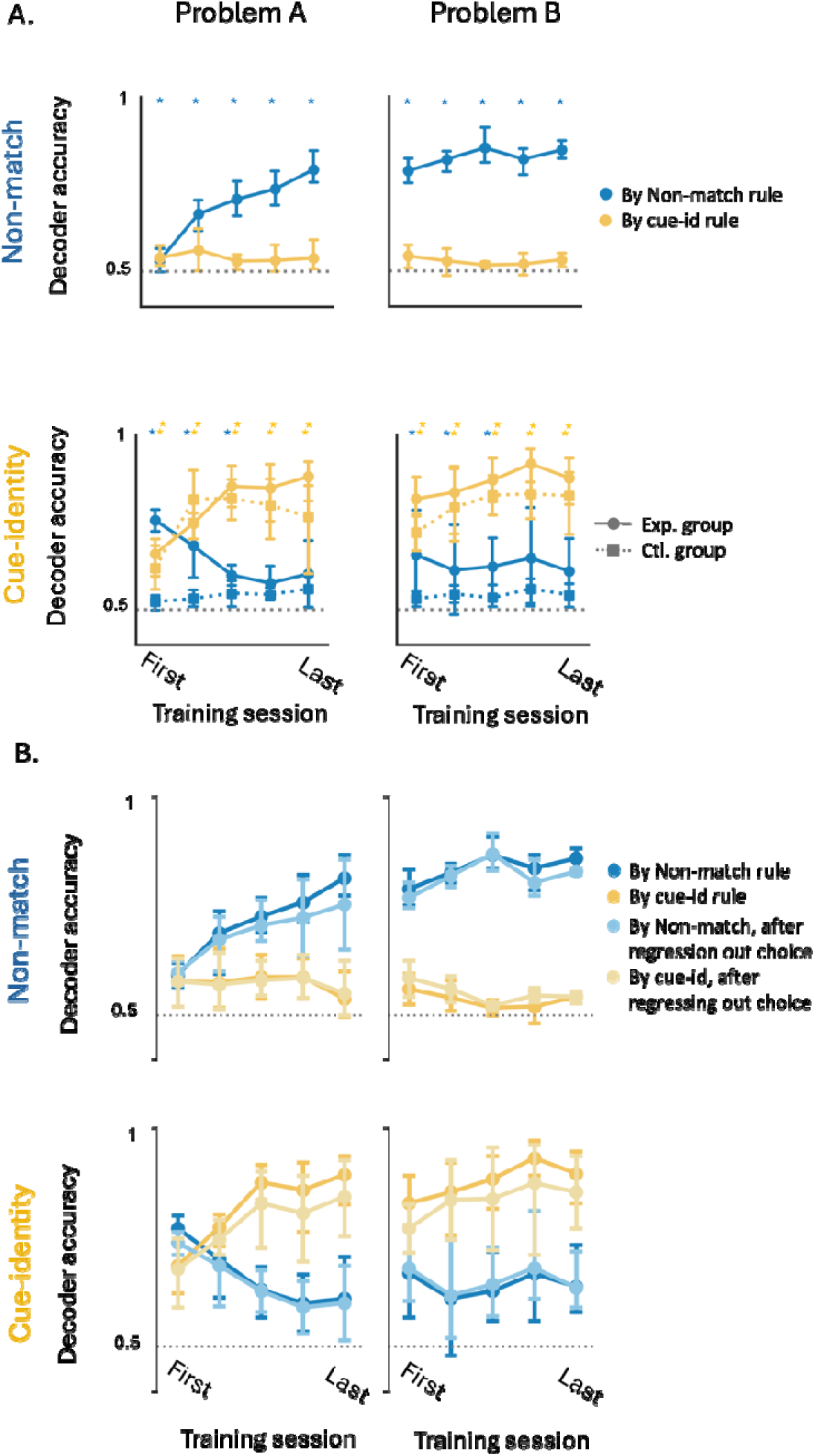
**A.** Neural decoder accuracy based on the non-match rule (blue) and cue-identity rule (yellow) in the main group of rats (circles) and the control group (squares, dashed lines). Asterisks indicate statistically significant differences from shuffled data for the non-match rule (blue) and cue-identity rule (orange) (p < 0.01, permutation test). Asterisks in the upper row denote statistically significant differences for the control group. **B.** Decoder accuracy based on the non-match rule (blue) and cue-identity rule (yellow), shown before (dark lines) and after (light lines) regressing out the rats’ choice (go/no-go).

